# An evolutionarily conserved scheme for reformatting odor concentration in early olfactory circuits

**DOI:** 10.1101/2025.01.23.634259

**Authors:** Yang Shen, Arkarup Banerjee, Dinu F. Albeanu, Saket Navlakha

**Affiliations:** Cold Spring Harbor Laboratory, Cold Spring Harbor, NY, USA

**Keywords:** olfactory circuits, odor concentration, divisive normalization, computational modeling, circuit evolution

## Abstract

Understanding how stimuli from the sensory periphery are progressively reformatted to yield useful representations is a fundamental challenge in neuroscience. In olfaction, assessing odor concentration is key for many behaviors, such as tracking and navigation. Initially, as odor concentration increases, the average response of first-order sensory neurons also increases. However, the average response of second-order neurons remains flat with increasing concentration – a transformation that is believed to help with concentration-invariant odor identification, but that seemingly discards concentration information before it could be sent to higher brain regions. By combining neural data analyses from diverse species with computational modeling, we propose strategies by which second-order neurons preserve concentration information, despite flat mean responses at the population level. We find that individual second-order neurons have diverse concentration response curves that are unique to each odorant — some neurons respond more with higher concentration and others respond less — and together this diversity generates distinct combinatorial representations for different concentrations. We show that this encoding scheme can be recapitulated using a circuit computation, called divisive normalization, and we derive sufficient conditions for this diversity to emerge. We then discuss two mechanisms (spike rate vs. timing based) by which higher order brain regions may decode odor concentration from the reformatted representations. Since vertebrate and invertebrate olfactory systems likely evolved independently, our findings suggest that evolution converged on similar algorithmic solutions despite stark differences in neural circuit architectures. Finally, in land vertebrates a parallel olfactory pathway has evolved whose second-order neurons do not exhibit such diverse response curves; rather neurons in this pathway represent concentration information in a more monotonic fashion on average, potentially allowing for easier odor localization and identification at the expense of increased energy use.

## Introduction

A central task of sensory systems is to reformat representations of sensory stimuli to efficiently encode key stimulus features. For example, in vision, brightness information captured by the retina needs to be preserved along the visual pathway to facilitate accurate perception of light intensity and contrast in the environment (1). Similarly, in audition, sound intensity captured by hair cells in the cochlea needs to be preserved to facilitate behaviors that require sound localization (2). While these transformations are well-studied for visual and auditory stimuli (3–7), much less is known about analogous transformations in the olfactory system. The ability to reliably estimate odor concentration (i.e., how much of an odor is present) is critical for olfactory navigation across species (8, 9). Here, we investigate the transformation of this key feature by early olfactory circuits.

Reliably tracking odor concentration is challenging (10). Based on receptor-ligand interactions, first-order (sensory) neurons increase their firing rates as odor concentration increases (11–15), such that the population activity of sensory neurons scales with odor concentration. However, this comes at a cost of odor discrimination, as representations become increasingly overlapping at high concentrations (16–19). In contrast, the average population activity of second-order and third-order neurons remains relatively flat over concentration changes that vary over many orders of magnitude (20– 22). This property is thought to be critical for concentration-invariant identification of an odorant (20, 23–25). However, this creates a conundrum: if concentration information is discarded in second-order neurons, which exclusively transmit odor information to the rest of the brain, how does the brain support olfactory behaviors, such as tracking and navigation?

By combining neural data analyses from diverse species with computational modeling, we provide a resolution to this issue and offer four contributions: First, we show how concentration information can be represented in second-order neurons, despite mean response flatness at the population level, and we show that this property is conserved across three species (locusts, zebrafish, and mice). Second, we demonstrate that a unitary circuit computation — divisive normalization — can generate all primary features of the reformatted second-order neuron representation given sufficient conditions on first-order responses. Third, we describe how this reformatted representation can be used to decode odor concentration using two biologically plausible strategies (rate-based and time-based decoding), and we discuss decoding constraints imposed by each scheme. Fourth, in land vertebrates, evolution has invented a parallel olfactory pathway whose response properties are much closer to those of first-order neurons; i.e., responses of second-order neurons in this pathway scale with concentration. We show that this representation substantially improves concentration decoding and odor localization (26), albeit with higher energy costs. Moreover, we hypothesize that the emergence of a parallel pathway in land vertebrates, in conjunction with the presence of long-range cortical feed-back loops, may have afforded neurons of the first pathway to carry out higher-order computations extending beyond simply representing sensory features.

Together, this study provides a unified theoretical framework for understanding how concentration information is reformatted by early olfactory circuits separated by several hundred-million years of evolution, from insects to mammals. Given the likely independent evolution of olfactory systems in vertebrates and invertebrates, our results suggest that evolution has converged to a common algorithm despite differences in the underlying circuit implementations.

## Results

Olfaction begins with olfactory sensory neurons (OSNs) that each expresses a single type of odorant receptor protein that binds with specific odor molecules (Fig. 1A). These first-order neurons transmit odor information to second-order neurons, such as projection neurons in insects, or mitral cells in vertebrates. The activity of second-order neurons is a combination of first-order neuron activity, as well as interneuron modifications (e.g., from local interactions in the antennal lobe of insects (27), or from local interactions in the olfactory bulb and long-range feedback in vertebrates (28–30)). Only second-order neurons project to the rest of the brain (e.g., the mushroom body in insects, the olfactory cortex in vertebrates), which subsequently orchestrate complex olfactory behaviors, such as odor discrimination and tracking (31). Hence, it is critical for second-order neurons to preserve key features of odors, including their identity and concentration. To determine if there is a conserved strategy of encoding odor concentration in olfactory circuits, we studied odor responses of first- and second-order neurons across several species. For first-order neurons, we re-analyzed data from fruit flies (24 odorant sensory neurons to 9 fruit odors across 4 concentrations (32), spanning 6 orders of magnitude in relative dilution) and mice (47 glomeruli to 5 odors across 4 concentrations (33), spanning 3 orders of magnitude). For second-order neurons, we re-analyzed data from locusts (110 projection neurons to 3 odors at 5 concentrations (21), spanning 3 orders of magnitude), mice (392 mitral cells to 5 odors at 4 concentrations (34), spanning 3 orders of magnitude) and zebrafish (358 mitral cells to 1 odor at 5 concentrations (22), spanning 4 orders of magnitude).

**Fig. 1.**
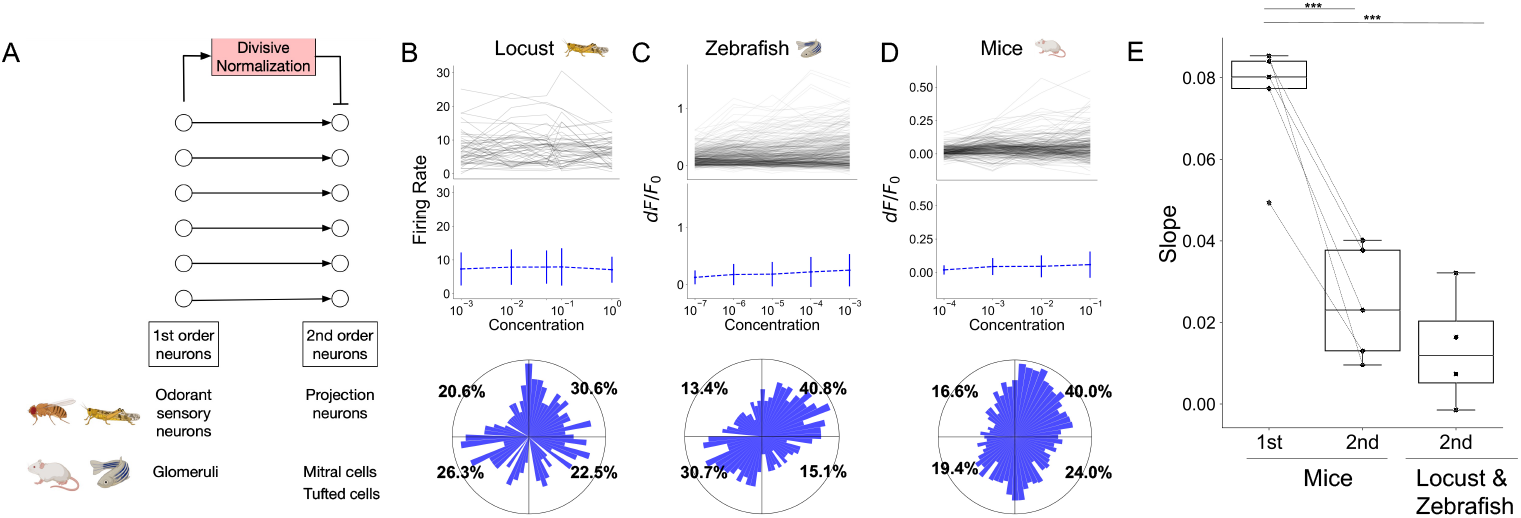
Second-order neurons across species demonstrate two common properties: flat average response of the population and diverse response shapes of individual neurons. **(A)** Illustration of the early olfactory circuit across species. **(B–D)** Example responses of second-order neurons in locust (to odor geraniol), zebrafish (phenylalanine), and mice (valeraldhyde), respectively. Top: individual neuron response curves across concentrations. Middle: mean population responses across concentrations. Bottom: polar histogram showing the distribution of individual neuron response shapes for all odors combined. The x-axis shows the slope between neuron response at concentration level 1 (lowest) to concentration level 3, and the y-axis shows the slope from concentration level 2 to concentration level 4 (highest). The angle of the bar shows the direction of the vector formed by these two slopes, and the length of the bar shows the percentage of neurons with the same angle. Top right quadrant: monotonically increasing responses; top left quadrant: decreasing then increasing; bottom left quadrant: monotonically decreasing; bottom right quadrant: increasing then decreasing. **(E)** Fitted slopes for mean population responses across concentrations for first- and second-order neurons. Each dot represents a single odor. First two columns show data collected in mice, and dashed lines connect the same odor. The third column shows all odors tested in locust and zebrafish combined. T-test is performed between the slopes formed by first- and second-order neurons; *** p-value < 0.001.

### Shared properties of odor concentration encoding in second-order neurons

We observed two common properties of odor representations in second-order neurons across all three species (Fig. 1B–D, top panels). First, for each odor, the average population response of second-order neurons remains nearly flat across concentrations that range 3–4 orders of magnitude. The fitted slopes of average responses of second-order neurons across increasing concentration levels are significantly lower than those of first-order neurons (Fig. 1E), consistent with previous results (21, 22, 33–36).

Second, the response curves of individual second-order neurons across increasing concentration levels have diverse shapes (Fig. 1B–D, bottom panels). We quantified the diversity of each neuron’s concentration response curve by computing the slopes between responses across concentration levels. The change in response across consecutive concentration levels may not be robust due to experimental noise and the somewhat limited range of concentrations sampled. Thus, for better illustration and ease of analysis, we computed the slopes between responses separated by 2 dilution levels, corresponding roughly to a 100x range in concentration. For example, the two slopes per neuron were computed using responses from concentration level 3 to level 1 and from level 4 to level 2. We defined the neuron to have a monotonically increasing (or decreasing) concentration response shape if both slopes are positive (negative). Non-monotonic concentration response shapes represent responses that increase and then decrease, or that decrease and then increase. We then calculated the angle between these two slopes for each neuron and plotted a polar histogram of these angles over all neurons (Fig. S1). Each quadrant of the polar histogram represents one of the four possible shapes (top right: monotonically increasing, top left: decrease then increase, bottom left: monotonically decreasing, bottom right: increase then decrease), and the percentage shown in each quadrant shows the percentage of neurons with the corresponding shape. We found diverse response shapes for second-order neurons in all three species (Fig. 1B–D). For example, for projection neurons in locust, 30.6% of neurons monotonically increased (top right quadrant), 26.3% monotonically decreased, 20.6% decreased then increased, and 22.5% increased then decreased. The response shapes were similarly spread over the four quadrants for mitral cells in both mice (40.0%, 19.4%, 16.6%, 24.0%) and zebrafish (40.8%, 30.7%, 13.4%, 15.1%). Thus, even though the average responses of second-order neurons are flat across concentrations, the response shapes of individual neurons are diverse.

In summary, two properties of second-order neuron responses across concentrations — flat average response of the population and diverse response shapes of individual neurons — are shared across locusts, zebrafish and mice.

### Divisive normalization recapitulates two ubiquitous properties of second-order olfactory neurons

How might such diverse shapes be generated from first-order neurons that respond more monotonically (37, 38)? One clue is the mean flatness of second-order responses, which hints at some type of normalization mechanism. We explored three commonly described mechanisms (22, 36, 39, 40) — divisive normalization (DN), subtractive normalization (SN), and intraglomerular gain control (IGC). Both divisive and subtractive normalization are global schemes based on population level activity, whereas intraglomerular gain control is a local normalization scheme where the activity of a neuron is normalized by only its own activity. We tested whether these normalization mechanisms could generate the two properties of second-order neurons described above (flat average response of the population and diverse response of individual neurons, across concentration levels) starting from experimentally observed first-order neuron responses in fruit flies and mice (Fig 2Ai–Aii).

**Fig. 2.**
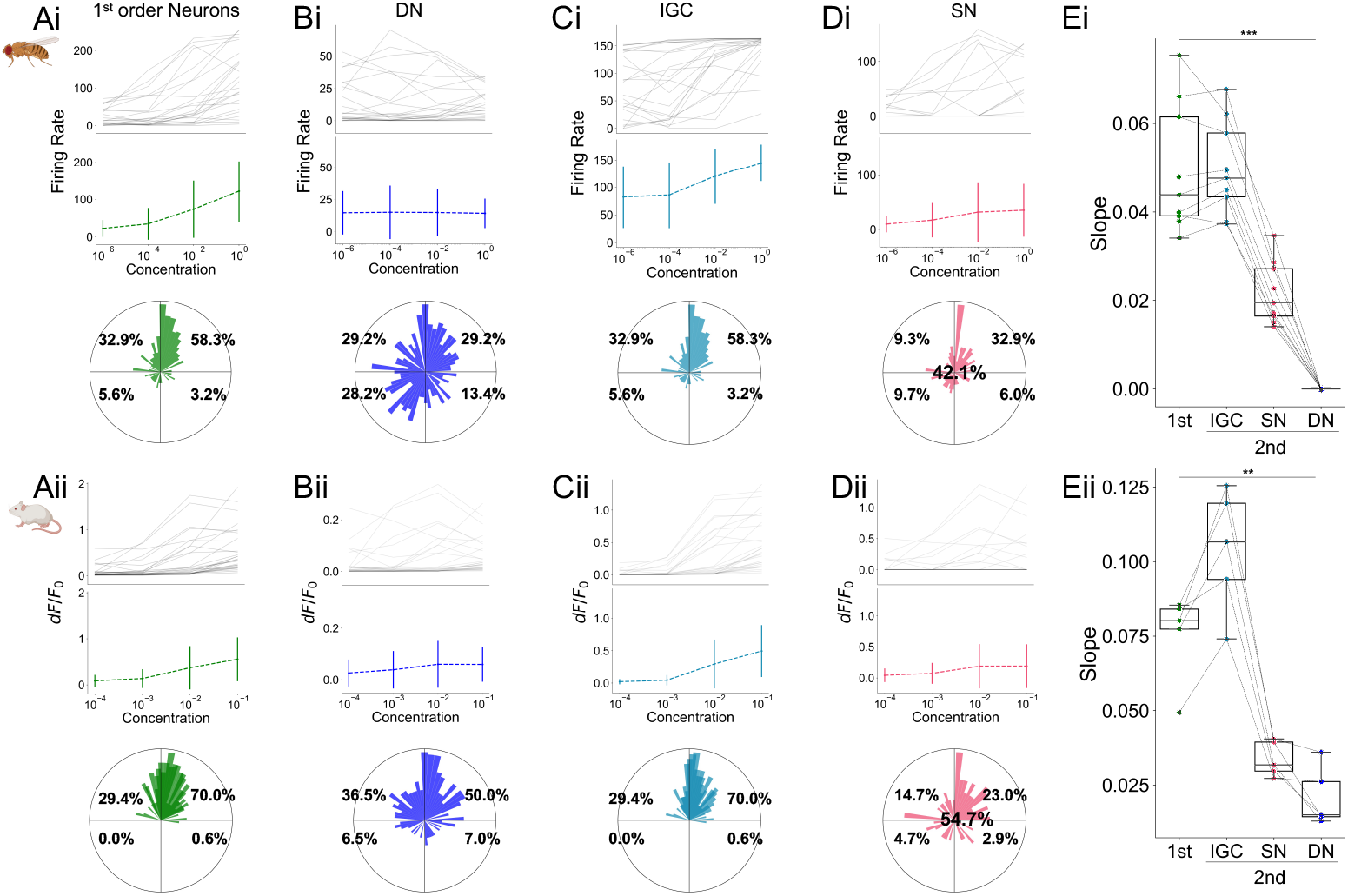
Responses of simulated second-order neurons recapitulate the two common properties. **(Ai–Aii)** Example experimentally measured responses of first-order neurons in fruit flies (apple odor) and mice (heptanal). **(Bi–Bii)** Simulated responses of corresponding second-order neurons after applying divisive normalization (DN) to first-order neurons. The top two rows show results of a single example odor, while the bottom row (the polar plots) show all odors combined. **(Ci–Cii)** Same as B but after applying subtractive normalization (SN). **(Di–Dii)** Same as B after applying intraglomerular gain control (IGC). **(Ei–Eii)** Fitted slopes for the mean population response across concentrations for first-order neurons, as well as simulated second-order neurons after IGC, SN, or DN is applied. Each dot represents an odor. Dashed lines connect the same odor across normalization methods. Paired t-test is performed between the slopes formed by first- and second-order neurons; *** p-value < 0.001, ** p-value < 0.01.

Divisive normalization divides the activity of a single neuron by the summed population activity (22, 36):

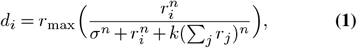

where *d*_*i*_ is the activity of the *i*^th^ second-order neuron, *r*_*i*_ is the activity of the corresponding *i*^th^ first-order neuron, *r*_max_ is the estimated maximum response of first-order neurons (determined individually for each dataset), *σ >* 0 controls the neuron’s sensitivity to normalization, *k* ∈ (0, 1) controls the strength of normalization, and *n >* 0 controls the shape of normalization curve. See Methods for parameter settings and equations for SN and IGC.

We found that after divisive normalization was applied to experimental first-order neuron responses, the simulated second-order neuron responses became significantly flatter over concentrations in both flies (1st order neuron slopes: 0.05±0.01 vs. 2nd order slopes: 0.000023±10^−4^; *p <* 0.001, Fig. 2Bi, Ei) and in mice (1st order: 0.08±0.01 vs. 2nd order: 0.02±0.01; *p <* 0.01, Fig. 2Bii, Eii). Thus, the average population responses of the simulated second-order neurons are nearly flat across concentrations, generating the first property. For the second property, we found that simulated second-order response shapes become more uniformly distributed in the polar plot histogram compared to those of first-order neurons. For example, the percentage of monotonically increasing neurons dropped from 58.3% in first-order neurons to 29.2% in second-order neurons in flies (Fig. 2Bi) and from 70.0% to 50.0% in mice (Fig. 2Bii). Thus, divisive normalization can reproduce both properties of second-order neurons.

In contrast, neither subtractive normalization nor intraglomerular gain control reproduce both properties of second-order neurons (Fig. 2C–E). Subtractive normalization recapitulates mean flatness at the population level in a relatively small parameter window (at the cost of silencing a large portion of neurons); however, it cannot reliably reproduce diverse response curves. Intraglomerular gain control fails to generate either property.

While divisive normalization has been previously shown to generate flat mean responses at the population level (22, 36), its role towards generating diverse concentration response curves has not been appreciated, and this latter property, as we will show, is critical towards decoding concentration levels downstream.

### Sufficient conditions for generating diverse concentration response curves via divisive normalization

Do these two properties inevitably arise as a result of divisive normalization applied to *any* first-order neuron responses, or do first-order responses need to have certain structure themselves? To answer this question, we synthesized concentration response curves for first-order neurons. These responses (*r*) to logarithmic scale concentration levels (*x*) follows the logistic function (37, 38):

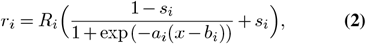

where *R*_*i*_ is the saturation level of neuron *i, a* controls the rate of saturation, *b* determines the location where the response of the neuron reaches half-maximum, and *s* is a small positive parameter representing the amount of spontaneous activity in the absence of stimuli.

We explored two classes of first-order neuron concentration responses. The first class does not exhibit cross-overs; i.e., the rank order of neurons, from the highest response amplitude to the lowest, remains the same across all concentrations. The second class of concentration responses exhibited cross-overs, which are closer to experimental first-order neuron responses due to combinatorial odor binding properties of OSNs (12–15, 32, 41). Here, the response amplitude ranks change, reflecting different rates of activity increase in neurons despite the overall trend of increasing. To simulate no cross-over responses (Fig. 3A), the values of *a, b*, and *s* are fixed for all neurons. To simulate cross-over responses (Fig. 3B), each simulated neuron has different values of *a, b*, and *s* sampled from independent uniform distributions. For both classes, the saturation level (*R*) is sampled from a Gamma distribution, which fits the experimentally observed activities of first-order neurons (Methods). By construction, for both classes, individual and average first-order neuron responses grow monotonically with increasing concentration level (Fig. 3A–B).

**Fig. 3.**
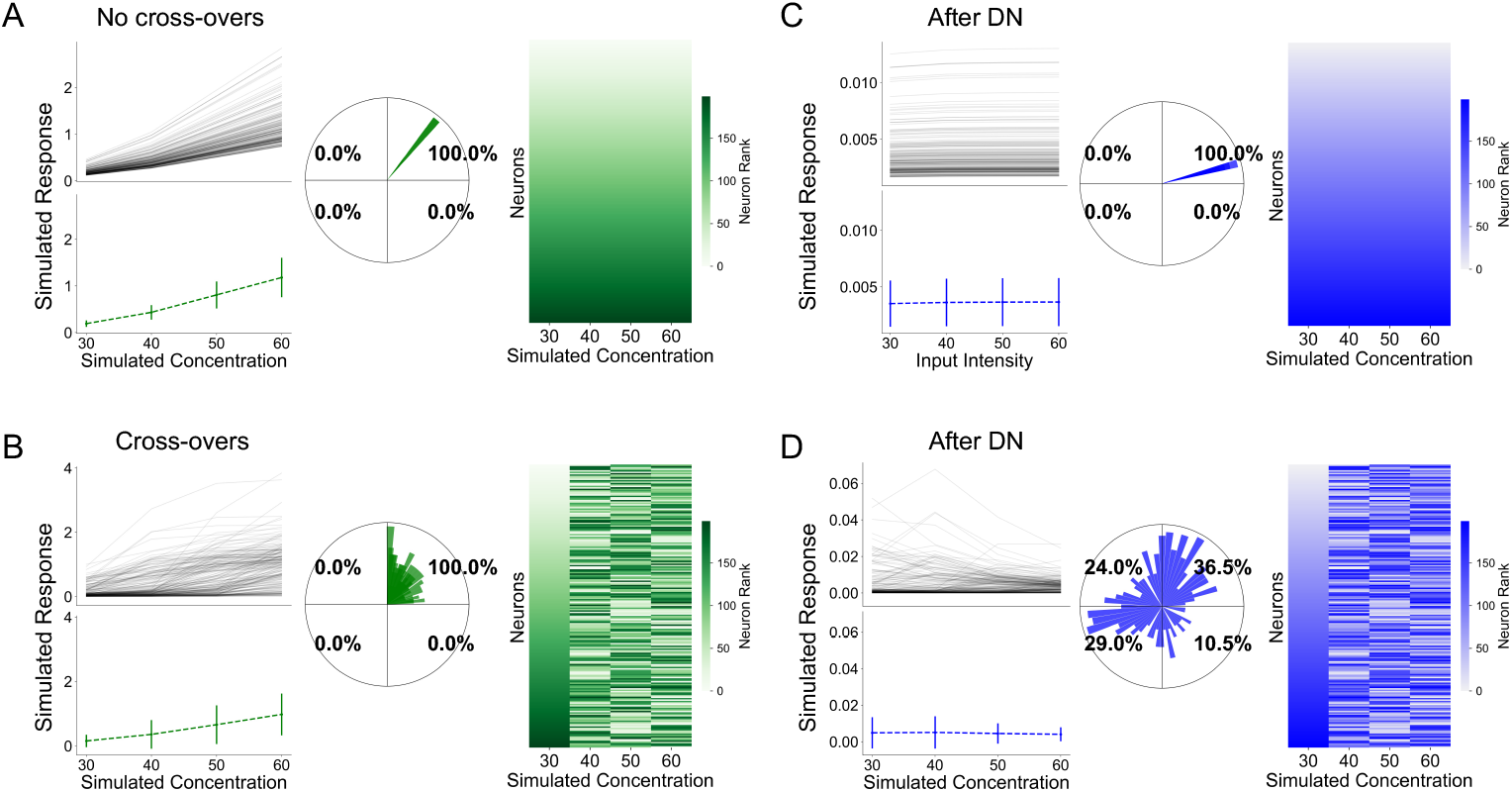
Sufficient conditions to generate diverse concentration response shapes using divisive normalization. **(A–B)** Simulated response curves of first-order neurons across concentration levels. Two classes of first-order responses are considered: those with cross-overs and those without cross-overs. Left: single neuron response curves and population average across four simulated concentration levels. Center: polar histogram for the shapes of response curves. Right: rank order of neuron activities at each concentration level (neurons are sorted according to response strength at the lowest nominal concentration, and the same indexing is preserved across the higher concentrations). **(C–D)**. Same as A and B but showing the responses of simulated second-order neurons after divisive normalization is applied to first-order neurons.

We found that the observed diversity of response curves in second-order neurons could only be reproduced after applying divisive normalization to first-order responses that had cross-overs, as opposed to no cross-overs (Fig. 3C–D). Specifically, for the former, the bars in the polar plot histogram of second-order neuron responses span all quadrants, whereas for no cross-over responses, the bars are exclusively located in the upper-right quadrant (i.e., monotonically increasing).

Why do diverse response curves emerge only from cross-over first-order responses? Divisive normalization (Eqn. Eq. (1)) effectively calculates the relative contribution of a neuron — i.e., its activity divided by the total activity of all neurons in the population – since the summation term *k*(*Σ*_*j*_*r*_*j*_)*^n^* dominates the 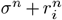 term in the denominator. Therefore, divisive normalization mostly does not alter the relative contribution (rank order) of each neuron in the ensemble. To create diverse shapes in second-order neurons, the first-order neurons must themselves increase their responsiveness with sufficient diversity (faster or slower, or steeper vs. shallower, compared to the average changes in activity of the population) such that their rank orders change across concentration levels. Biologically, as concentration increases, cross-overs emerge as a consequence of competing growth rates of individual first-order neurons versus the population. In summary, cross-over first-order responses are sufficient for divisive normalization to generate non-monotonic responses in second-order neurons.

### Two complementary schemes of decoding odor concentration from populations of second-order neurons

How can higher order brain regions decode concentration information from second-order neurons that have flat mean responses, but diverse individual concentration response curves? We discuss two potential strategies, one using firing rate information and the other using spike timing information.

First, we explored how concentration information can be decoded from the diverse shapes of response curves of second-order neurons. We used a simple multi-class logistic regression model to predict the concentration of an odor using the experimental odorant response activities of second-order neurons, where the model was trained using all but one trial and tested on the held-out trial (Methods). We found that accuracies were nearly-perfect in all three species (Fig. 4A): 100±0% in locusts, 96±4% in zebrafish, and 97±4% in mice (chance accuracy is 25% for data with four concentration levels, and 20% for five concentration levels). Moreover, this decoding accuracy is on par with that derived using first-order neuron responses (Fig. 4B) and corresponding simulated responses of second-order neurons after divisive normalization is applied (Fig. 4C). Hence, divisive normalization reformats concentration information, so that it is no longer coded by the amplitude of mean population responses, but by the diverse shapes of response curves. In other words, each concentration level has its own combinatorial code in second-order neurons, allowing for the concentration to be decoded.

**Fig. 4.**
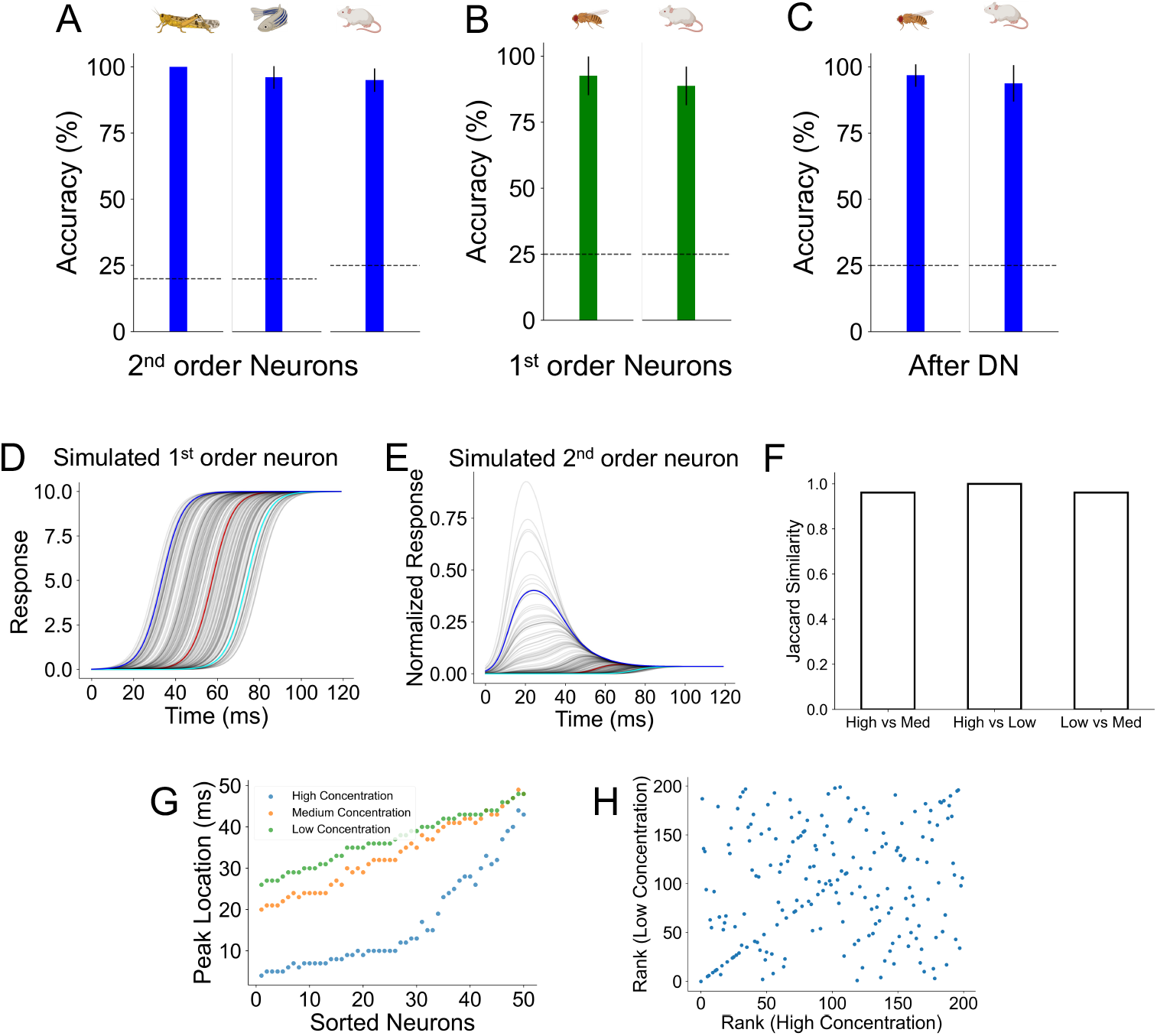
Divisive normalization reformats concentration representations in the rate and time domains. **(A)** Accuracy for concentration classification using experimental responses of second-order neurons (left: locust; center: zebrafish; right: mice). The concentration levels of each odor are classified independently per odor. Reported are the average accuracy across all odors, and error bars show standard deviations. Dashed line shows the accuracy level for random guess (0.2 for locust and zebrafish, 0.25 for mice). **(B)** Same as A but for experimental first-order neurons in fruit flies and mice. **(C)** Same as A but after divisive normalization applied to first-order responses in B. **(D)** The temporal response curves of simulated first-order neurons to an odor. Neurons with different affinity to this synthetic odor are highlighted (high affinity: blue; medium affinity: red; low affinity: cyan). **(E)** Same as D, after divisive normalization is applied. **(F)** Jaccard similarity between the sets of the 50 earliest-responding neurons for each pair of concentration levels. The value of Jaccard similarity ranges between 0 and 1; 1 means the two sets of neurons contain identical neurons, and 0 means they share no neurons in common. **(G)** The time when the 50 earliest-responding neurons respond under each concentration level. The neurons are sorted by their peak time under low concentration. **(H)** Rank of neurons under low concentration based on their time of peak response (y-axis) vs. rank of neurons under high concentration (x-axis).

Second, we considered a complementary scheme by which odor concentration can be decoded using information in the time-domain. Latency or primacy coding (42–44) proposes a mechanism to achieve concentration-invariant odor recognition by using information in only a few second-order neurons that have the highest affinity to an odor, i.e., neurons that respond early and stably to the odor across concentrations. To determine if this mechanism could in addition decode concentration levels, we simulated the temporal responses of 200 first-order neurons to three concentration levels of an odor using logistic functions with different parameter values; e.g., one parameter value controls the sensitivity of the response curve (Fig. 4D; Methods). After divisive normalization, the logistic response curves of first-order neurons transformed into non-monotonic curves (Fig. 4E). This agrees with the temporal response shapes of second-order neurons observed experimentally (45, 46).

We report three observations from these simulations. First, the set of second-order neurons with the highest-affinities (where affinity is based on the time latency of response) remained almost the same across concentrations levels: the Jaccard similarity between the sets of highest-affinity neurons for each pair of concentration levels was *>* 0.96, Fig. 4F). Second, the average timing of the peak response (i.e., the response latency) of these primacy neurons, decreased as concentration increases (Fig. 4G). This is not surprising and is simply meant to model the binding of odorants to odorant receptors with its resulting inverse correlation between response amplitude and latency (higher amplitude → shorter latency) (47). Third, critically, the rank orders of the set of primacy neurons were scrambled across concentrations, due to cross-over responses of corresponding first-order neurons. For the 10 and 50 highest-affinity neurons responding to the high concentration level, 7 (70%) and 32 (64%) shifted ranks, respectively, compared to their corresponding ranks at low concentration; for all 200 simulated neurons, 164 (82%) shifted ranks from high to low concentration (Fig. 4H).

These observations have major implications for both concentration-invariant odor identification and odor concentration decoding by third-order neurons under the primacy model. Odor identity can be encoded by the set of highest-affinity neurons (which remains invariant across concentrations); however, the downstream read-out mechanism must not be sensitive to their rank order. Odor concentration can be encoded using either the average response latency of neurons (which scales with concentration), or alternatively, by learning to associate different temporal sequences of neuronal responses with different concentrations.

Hence, two complementary strategies emerge in decoding odor concentration by higher-order brain regions. Divisive normalization, operating in antennal lobe (insects) or olfactory bulb (mammals) circuits, is a potential mechanism for generating odor-specific subsets of second-order neurons whose combinatorial activity or whose response latencies represents concentration information. Importantly, in both schemes, some form of learning within downstream neural circuits is required.

### A parallel pathway for odor representation in land vertebrates better supports odor localization

Locating an odor source is essential for survival in the wild. While olfactory navigation over large distances can be solved by serial sniffing (48), at close distances rodents rely more on stereo olfaction to localize stimuli (49–52). This requires that the brain quickly determine: 1) if there exists a difference in odor concentrations between two sniffs or between signals from two nostrils in a single sniff; 2) if there is a difference, which sniff/nostril sensed a higher concentration; and 3) how strong the difference is.

We showed above that odor concentration could be decoded from second-order neurons, but the proposed mechanisms require additional learning in downstream neural circuits, which may be slow. Odor localization may be achieved more easily and quickly (e.g., by a simple comparator across nostrils) if a direct copy of concentration information, where neuronal responses increase monotonically with concentration, could be stored beyond first-order neurons.

Indeed, evolution has discovered a parallel pathway in land vertebrates involving an additional type of second-order neuron — the tufted cells. Compared to mitral cells, tufted cells on average respond faster and more reliably to lower concentrations of an odorant, and their activity increases nearly monotonically with increasing odor concentration (19, 26, 53–57) (Fig. 5A–B), resembling first-order neuron (glomerular) activity. Previous work showed that tufted cell ensembles outperform mitral cell ensembles in concentration classification and odor generalization tasks (26).

**Fig. 5.**
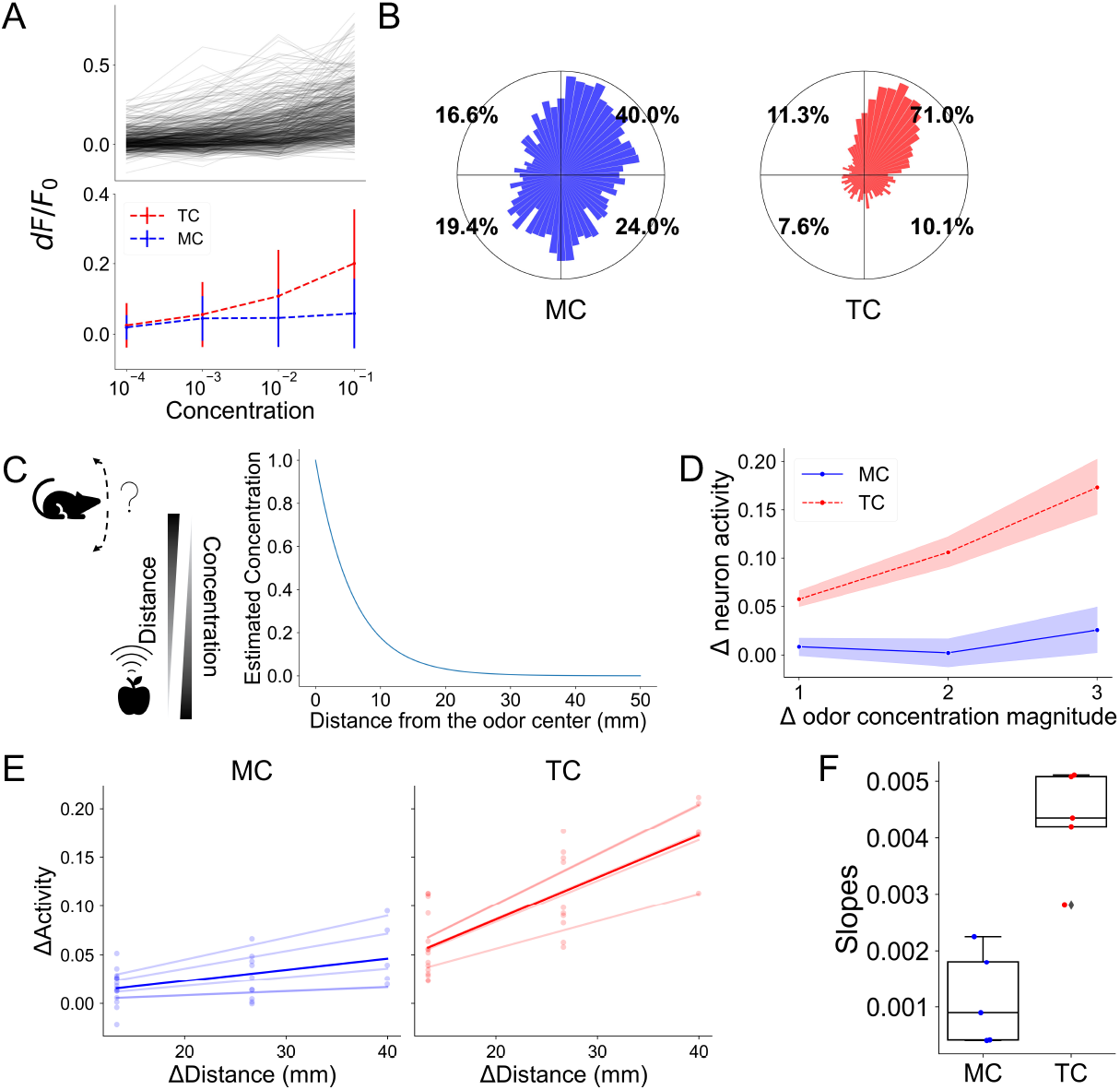
A parallel olfactory pathway in land vertebrates provides improved concentration decoding. **(A)** Top: concentration response curves of individual tufted cells to a single odor (valeraldhyde). Bottom: average population responses for mitral (MC, blue) and tufted (TC, red) cells. **(B)** Polar histogram showing the distribution of concentration response shapes MCs and TCs for all five odors combined. **(C)** Left: Cartoon illustration for the odor localization task. Right: estimated decrease in odor concentration as distance from the odor source increases. **(D)** Change in average population neuron activities across all pairs of concentrations. The x-axis shows the absolute value of difference in the relative dilution (in order of magnitude) between the two concentration levels. The y-axis shows the change in activity. Results are averaged over five odors, and the shaded area shows the 95% confidence intervals. **(E)** Estimated change of neural activities as the distance between the odor source and the animal’s location increases for mitral cells (left) and tufted cells (right). Each dot shows a particular concentration for an odor. There are 20 dots in total (4 concentrations x 5 odors). Each opaque line is a linear regression calculated based on all concentrations of a given odor, and the darker lines are the average over all odors. **(F)** Slopes for the linear regression lines shown in E. Each dot represents an odor. The value of the slope is an estimate of neural sensitivity to the change in distance to the odor source. The higher the slope, the higher the sensitivity.

Here we show that in addition to these tasks, tufted cells are better suited to solve the stereo odor localization problem (Fig. 5C, left). We re-analyzed second-order neurons in mice (34), consisting of 392 mitral cell and 387 tufted cell responses to 5 odors across 4 concentration levels that span 3 orders of magnitude in relative dilution. We found that the same concentration change causes a significantly larger change in tufted cell responses compared to mitral cells (Fig. 5D). This predicts that tufted cells should better discriminate smaller concentration changes. Relying on experimentally measured fall-off of concentration with distance from an odor source (Fig. 5C, right), we indeed find that the tufted cells are better at resolving smaller distances — the same increase in the distance from the odor center would lead to larger change in tufted cell activity compared to mitral cells (Fig. 5E–F). However, this increased performance comes at a cost; the average population activity level of tufted cells is roughly three times higher than in mitral cells (0.03±0.01 for mitral cells vs 0.09±0.03 for tufted cells), indicating that tufted cells consume more energy than mitral cells. Thus, we posit that the evolution of the tufted cell output channel traded-off energy for faster and finer concentration discrimination, which is essential for odor tracking and navigation in land vertebrates.

## Discussion

### Summary of findings

Differentiating odor concentration is critical for olfactory behaviors, including odor-based navigation and localization. First-order olfactory neurons exhibit increasing responses to higher levels of concentration, but second-order neurons on average remain flat across many orders of magnitude of concentration fluctuations. Such concentration invariance is thought to be beneficial for identifying odorants independent of concentration (20, 25). However, under this model, it has remained unclear how odor concentration information can be readily accessed by higher brain regions to guide behavior and decision-making. Our analysis, based on previous experimental work (22, 33, 35, 36), shows that circuits in the early olfactory system across species (locusts, zebrafish, and mice) normalize overall activity of second-order neurons, while also retaining adequate diversity in the individual second-order neuron concentration response curves: some neurons increase or decrease their activity monotonically with concentration, whereas others respond non-monotonically. This diversity enables individual concentration levels to be encoded combinatorially, which can then be decoded using spike rates or spike times. Moreover, we analyzed an additional type of second-order neuron (tufted cells) that has evolved in land vertebrates and that outperforms mitral cells in concentration encoding (26) and odor localization tasks. Indeed, recent work suggests that the anterior olfactory nucleus, which gets stronger input from tufted cells than mitral cells, plays a central role in decoding odor concentration (26, 58). These results suggest a trade-off between concentration decoding and normalization processes, which prevent saturation and reduce energy consumption.

What might be the benefits of evolving a parallel second-order output channel (tufted cells) in land vertebrates? We speculate that this alternative pathway may have enabled mitral cells to evolve new functionalities. By combining sensory information from first-order neurons with feedback from higher brain regions, mitral cells may be ideally suited to perform learning-dependent computations beyond simply relaying sensory input, such as predictive processing or integrating context within decision-making (59). It is also possible that the brain uses multiple concentration decoding strategies in parallel, depending on the complexity of the olfactory scene and the behavioral needs of the moment. Overall, our findings highlight the importance of an evolutionarily conserved computation in odor coding and how the key feature of odors is efficiently preserved along the olfactory pathway.

### Mechanisms for concentration decoding

Both concentration decoding schemes proposed here (spike-based and time-based) require learning in downstream circuitry — i.e., third-order neurons need to learn which temporal sequences of neurons are to be associated with each concentration level of an odor. This sequence may be further altered by noise present in natural environments, which may lead to additional variability across repeated observations of the same odorant concentration. Experimental results indeed show that while some neurons remain rank-stable, different subsets of early responding secondary neurons are silenced or start firing at higher vs. lower concentrations of the same odor (19, 26, 42, 60). As such, a substantial fraction of second-order neurons change rank depending on the concentration range and odor identity. In addition, the spiking temporal integration window of third-order neurons in downstream brain regions ought to be small enough to read-out the temporal shifts in response latencies across concentrations for primacy neurons. The complementary combinatorial response (firing rate) scheme also necessitates learning: different third-order neurons need to be associated, presumably via synaptic plasticity, with different concentrations of a given odorant. The feasibility of the learning algorithms for both decoding schemes remain to be tested experimentally.

Our work does not rule out alternative mechanisms for encoding concentration in olfactory circuits. For example, concentration can be encoded via the total activity in the trigeminal nerve, through the different latencies between the medial and lateral olfactory bulb under different concentrations (61), or via temporal integration via serial sniffs (counting the total number of odor packets over multiple sniffs) (48, 62).

### Experimental predictions

Our work raises two testable experimental predictions about the effects of disrupting circuitry involved in implementing divisive normalization on downstream encoding and behavior. First, the properties of second-order neurons, such as mean-flatness and diverse concentration response shapes enabled by divisive normalization, would disappear if the neural circuitry supporting divisive normalization is suppressed. In insects, experimental evidence suggests that divisive normalization is likely achieved through the network of inhibitory interneurons in the antennal lobe (63). In vertebrates, divisive normalization is likely implemented by short-axon cells (GABAergic-dopaminergic inhibitory cells in the olfactory bulb), whose axons provide interglomerular crosstalk (22, 33, 64–68). Indeed, three properties of short-axon cells in the olfactory bulb make them ideally suited to implement divisive normalization: odor responses of short-axon cells scale monotonically with concentration, they functionally inhibit mitral cells, and crucially, ablating these neurons shift the concentration response curves of mitral cells from being diverse to becoming more monotonic (33). Concomitantly, this loss of diverse response shapes in second-order neurons may hinder fine concentration discrimination of the same odor. Second, for mammals, prior work has hypothesized that concentration-invariant encoding requires recurrent processing in the piriform cortex (20, 44). In contrast, our results predict that concentration invariant neural responses, the diversity in individual neuron concentration responses, and the proposed latency/primacy representations for odor identification emerge due to computations within the olfactory bulb. Consequently, we predict that these properties will persist even after inactivating the cortex.

### Evolutionary conserved transformation of concentration encoding

Is there a shared computation employed by early olfactory circuits from flies to mammals to generate our proposed concentration encoding scheme? Our results indicate that divisive normalization is sufficient to explain two experimentally observed properties of second-order neurons: flat mean responses and diverse individual concentration response curves. As noted above, experimental evidence suggests that divisive normalization is implemented by distinct inhibitory circuits in the antennal lobe in insects vs. the olfactory bulb in mammals. Thus, while the circuit implementation of this transformation may be different across species that diverged over 100 million years ago, invoking Marr, the algorithmic description appears to be conserved. Given that the invertebrate olfactory receptors have evolved independently from those of vertebrates, our results also imply a functional convergence shaped by similar computational goals. Finally, in machine learning, applying divisive normalization to early encoding layers generates efficient coding representations (69, 70) that accelerate network training (71), revealing the broad applicability of this transformation in both biology and engineering.

## ACKNOWLEDGEMENTS

The authors are grateful to Zane Aldworth, Rainer Friedrich, and Elissa Hallem for sharing experimental data, and to members of the Banerjee, Albeanu, and Navlakha labs for constructive comments on the manuscript.

## Methods

### Analysis of experimental data

#### Mice

The experimental data was published in Chae et al. (26) (2022). Briefly, head-fixed, naïve mice were presented with 5 odors at 4 concentration levels (across three orders of magnitude), and two-photon calcium imaging was performed to measure responses from 392 mitral cells and 387 tufted cells. We also re-analyzed a dataset consisting of neuronal responses from 47 glomeruli to the same odor panel (33). Each measurement was repeated for 4-6 trials for each odor at each concentration. For each measurement, a monomolecular odor was presented for 4s, preceded by 10-12s of air baseline, followed by 7-10s air recovery. Responses (d*f* /*f*) were calculated based on the air baseline period, which was determined for each neuron independently. Neuronal responses for each cell-odor pair were defined as the average d*f* /*f* over the entire odor period. Data from all field of views and all odors were combined in our analysis.

#### Zebrafish

The zebrafish data was published by Zhu et al. (22) (2013). We re-analyzed calcium imaging responses of 358 mitral cells to a single odor across 5 concentration levels. Mean neuron responses were defined as the d*f* /*f* for the period after odor onset (about 2*s*).

#### Locust

The locust data was published by Stopfer et al. (21) (2003). We re-analyzed spiking activity of 110 projection neurons in response to 3 odors across 5 concentration levels per odor. Each experiment was repeated for 15 trials. Each trial began with a 2s air baseline period, followed by odor presentation for 1s.

#### Fruit fly

The Drosophila data was published by Hallem and Carlson (32) (2006). Spiking activity of 24 ORNs to 9 fruity odors across 4 concentrations were measured. Each ORN measurement was made in a different animal. All analyses were based on average responses of single neurons across the 6 trials (as shown in Table S2 of Hallem and Carlson (32)), except for the classification task of concentrations (Fig. 4B-C), in which the 6 trials were used individually.

#### Selecting responding neurons

For a given odor and given concentration, a significant neuronal response was determined by activity 2 standard deviations higher than the baseline rate for at least 20% of the trials (e.g., at least 3 trials out of total 15 trials, or 100% of the trials if there is only one trial). Additionally, if a neuron responds to at least one concentration for a given odor, the neuron is considered to be responding to the given odor. Only responding neurons are used for analysis.

### Normalization methods

#### Divisive Normalization (DN)

Divisive normalization is performed according to the equation below:

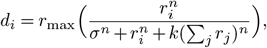

where *d*_*i*_ is the activity of the *i*^th^ second-order neuron, *r*_*i*_ is the activity of the corresponding *i*^th^ first-order neuron, *r*_max_ is the estimated maximum response of first-order neurons (determined individually for each dataset), *k* is a parameter controlling the strength of normalization, and *n* is a parameter controlling the shape of normalization curve. We fix *n* = 1.5 following Olsen et al. (36), and we fix *k* = 0.1, though we find no qualitative differences in our conclusions using different values of k (Fig. S2). Similarly, we use *σ* = 12 for the Drosophila data following Olsen et al. (36), and *σ* = 1 for all other datasets. This model of divisive normalization was modified from Olsen et al. (36).

#### Intraglomerular Gain Control (IGC)

Intraglomerular transformation modulates the responsiveness of a neuron based only on its own response as:

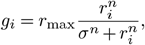

where *r*_*i*_ is the activity of the *i*^th^ first-order neuron, and *g*_*i*_ is the normalized activity of the corresponding second-order neuron. All other constants are fixed as above.

#### Subtractive Normalization (SN)

In subtractive normalization, each second-order neuron receives inhibition from interneurons proportional to the sum of first-order neuron activities:

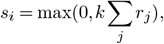

where *k* is the parameter controlling the strength of normalization, *r*_*i*_ is the activity of the *i*^th^ first-order neuron, and *s*_*i*_ is the normalized activity of the corresponding second-order neuron. We fix *k* = 1*/N* for subtractive normalization, where *N* is the number of first-order neurons.

#### Fitting slopes for mean population responses

The slopes of population mean responses (Fig. 1E, Fig. 2Ei–Eii) are fitted in a log-linear manner using LinearRegression function in the `sklearn` library of Python. The slope of each odor was fitted independently. Before fitting, the mean values were first min-max normalized by the minimum and maximum values of individual neurons.

#### OSN simulation

The 200 OSN curves are simulated using a logistic function:

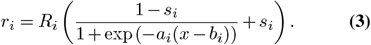

The parameter *R*_*i*_ (one per neuron) is sampled from a gamma distribution Γ(*α, β*), where *α* = 1.15, *β* = 1.92, parameters fitted from experimental data in mice glomeruli (). The *a* parameter controls how fast the curve reaches saturation value *R* (higher values of *a* indicate faster saturation but slower activity initiation). The *b* parameter controls the *x* value where the OSN activity reaches half-maximum of *r*_max_. The *s* parameter is a small non-zero parameter representing the amount of spontaneous activity in the absence of stimuli.

#### Concentration-response curves

When simulating concentration-response curves, the parameter *x* in Eqn. 3 is set to discrete concentration levels at (arbitrary) values of 30, 40, 50 and 60.

#### Non-crossover responses

The values of parameters *a, b*, and *s* are fixed at *a* = 0.1, *b* = 50, *s* = 0.

#### Crossover responses

The values of *a, b* and *s* are sampled from uniform distributions between [0.05, 0.4], [30, 80] and [0, 0.05] respectively.

#### Temporal response curves

When simulating temporal-response curves, the parameter *x* in Eqn. Eq. (3) indicates the time ranging from 0 to 120 ms. The parameters for different concentration levels are:

| Concentration level | $r_{max}$ | $a$ | $b$ | $s$ |
| --- | --- | --- | --- | --- |
| Low | 5 | 0.3 | Uniform[30,80] | 0 |
| Medium | 10 | 0.2 | Uniform[30,80] | 0 |
| High | 15 | 0.1 | Uniform[30,80] | 0 |

#### Logistic regression on concentration decoding

Multi-class logistic regressions were performed using the LogisticRe-gression function in the `sklearn` library of Python, with `liblinear` kernel and *f*_2_ regularization, and under the “one vs. rest” scheme.

For mice data, concentrations were first classified within each odor independently, and the reported accuracy was averaged over the five odors. There are four trials per given odor-concentration pair for glomeruli data, and five trials for mitral cell data; the regression model was trained on three/four trials randomly selected and tested on the left-out trial. This process was repeated ten times (with a different, random trial left out per concentration level each time), and the final accuracy for this odor is the average over the ten repeats. The concentrations in locust and fruit fly data were classified in the same way as mice data.

For zebrafish data, since there is only one measurement for each odor-concentration pair, trials for each concentration level were simulated by adding noise to the experimental data. In total 10 repeats were simulated for each concentration level. Noise was generated from a Gaussian distribution with mean equal to 0 and standard deviation equal to the mean response of all neurons in the given concentration.

#### Estimating distance based on odor concentration

The estimates of concentration fall-off as a function of distance from the odor source were based on experimental measurements (Albeanu lab, unpublished data). The mean odor intensity was measured using a photoionization detector (Aurora scientific) while laterally displacing the odor source from the PID in increments of 5 mm, over a total range of 50 mm, across 3 different odors. The observed relationship was well fit by the equation:

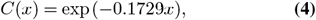

where *C*(*x*) is the normalized concentration at distance *x*, and *x* is the distance (in mm) between the location of measurement and the source of odor.

