## Supplement Information for "An evolutionarily conserved scheme for reformatting odor concentration in early olfactory circuits"

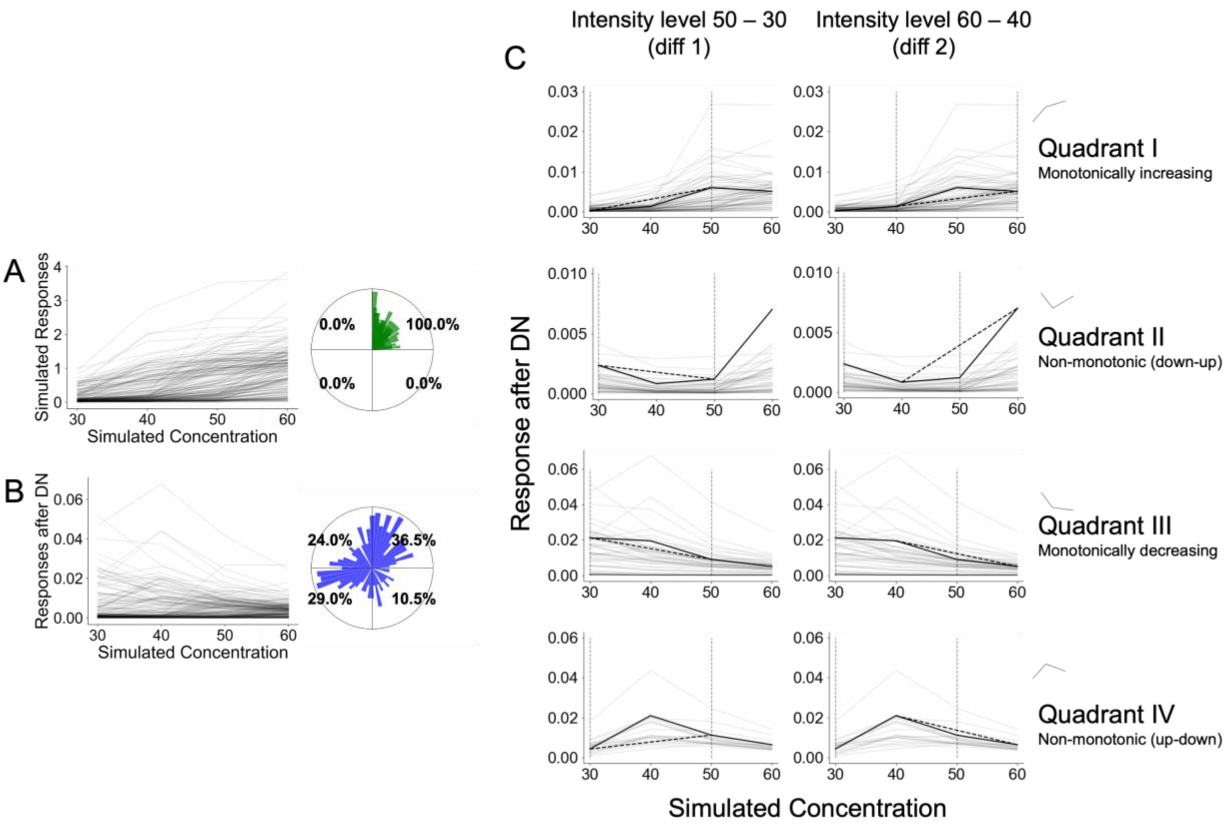


**Figure S1. Quantification of diverse concentration response shapes. (A)** Response curves of 200 simulated first-order neurons across four intensity levels (left) and the corresponding polar histogram (right). **(B)** Response curves and polar histogram of 200 simulated second-order neurons (after divisive normalization is applied to first-order neuron responses). **(C)** Response curves of single second-order neurons that belongs to each quadrant. The x and y axes in the subplots are the same as panel A. The four rows correspond to the four quadrants of the polar histogram. The two columns show the difference between single neuron responses at intensities 50 and 30 (diff1), and at intensities 60 and 40 (diff2). The four quadrants correspond to the four possible shapes of curves: top row (quadrant I): the response of a single neuron increased from intensity level 30 to 50, as well as from intensity level 40 to 60 (i.e., the angle formed by vector [diff1, diff2] is between 0 and $\pi/2$); second row (quadrant II): the response decreased from intensity level 30 to 50, but increased from intensity level 40 to 60 (i.e., the angle is between $\pi/2$ and $\pi$); third row (quadrant III): the response decreased from intensity level 30 to 50, as well as from intensity level 40 to 60 (i.e., the angle is between $\pi$ and $3\pi/2$); and fourth row (quadrant IV): the response decreased from intensity level 30 to 50, but increased from intensity level 40 to 60 (i.e., angle is between $3\pi/2$ and $2\pi$). Polar plots in A and B are normalized histograms (sum of the area of all bars equals 1). Each bar has the same angular width. The orientation of each bar shows the angle formed by [diff1, diff2], and the radial length shows the percentage of neurons that formed an angle within the angular width of the bar.

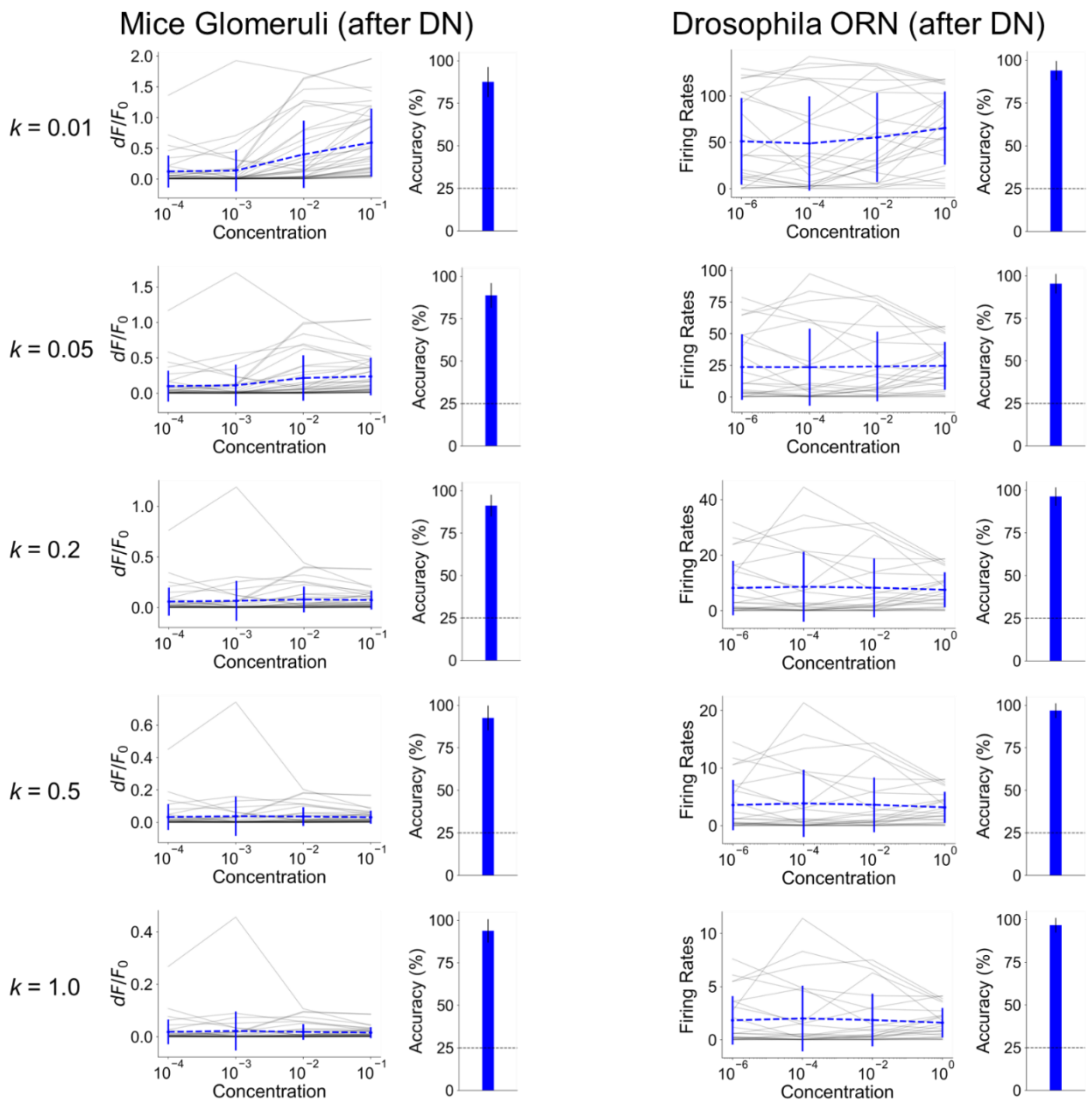


**Figure S2: Different *k* values have qualitatively similar effect on second-order neuron responses and concentration classification.** Divisive normalization applied to mice glomeruli (left) and Drosophila OSN (right) data under different values of parameter *k*. The mean population responses are generally flattened after divisive normalization, and the effect strengthens with increasing *k* values. In addition, the accuracy of concentration classification (the classification test is the same as Fig 4A–C) stays high under all *k* values.
